# A Hox code defines spinocerebellar neuron subtype regionalisation

**DOI:** 10.1101/640243

**Authors:** Eamon Coughlan, Victoria Garside, Siew Fen Lisa Wong, Huazheng Liang, Dominik Kraus, Kajari Karmakar, Upasana Maheshwari, Filippo M. Rijli, James Bourne, Edwina McGlinn

**Affiliations:** EMBL Australia, Australian Regenerative Medicine Institute, Monash University, Clayton, Vic, 3800, Australia; Neuroscience Research Australia, Randwick, NSW, 2031, Australia; Friedrich Miescher Institute for Biomedical Research, Maulbeerstrasse 66, 4058 Basel, Switzerland; University of Basel, 4001 Basel, Switzerland

**Keywords:** Hox gene, Hox cluster, spinocerebellar, proprioception, sensory neuron, miR-196, Hoxc9, microRNA, Gdnf

## Abstract

Coordinated body movement requires the integration of many sensory inputs. This includes proprioception, the sense of relative body position and force associated with movement. Proprioceptive information is relayed to the cerebellum via spinocerebellar neurons, located in the spinal cord within a number of major neuronal columns or as various scattered cell populations. Despite the importance of proprioception to fluid movement, a molecular understanding of spinocerebellar relay interneurons is only beginning to be explored, with limited knowledge of molecular heterogeneity within and between columns. Using fluorescent reporter knock-in mice, neuronal tracing and *in situ* hybridisation, we identify widespread expression of *Hox* cluster genes, including both protein-coding genes and microRNAs, within spinocerebellar neurons. We reveal a “*Hox* code” based on axial level and individual spinocerebellar column, which, at cervico-thoracic levels, is essential for subtype regionalisation. Specifically, we show that Hoxc9 function is required in most, but not all, cells of the major thoracic spinocerebellar column, Clarke’s column, revealing heterogeneity reliant on Hox signatures.

## Introduction

The human body navigates movement with astonishing success, displaying locomotor actions that are fluid and coordinated. Proprioceptive sensory neurons (PSNs), various subclasses of which are located within muscles, tendons and joints, convey information to the central nervous system regarding changes in muscle length and muscle force (Proske and Gandevia, 2012). This input, along with the perception of skin deformation sensed by cutaneous receptors (Shambes et al., 1978), provides a constant awareness of the relative positioning of body components such as the limbs. In humans, proprioception declines with age (Goble et al., 2009) and in numerous disease states (Borchers et al., 2013; Conte et al., 2013), manifesting as irregular movement trajectory, exaggerated force and postural instability (Goble et al., 2009). In mouse, the ability to selectively ablate PSNs has revealed their essential contribution to locomotor pattern generation (Akay et al., 2014), maintenance of spinal alignment (Blecher et al., 2017a) and the realignment of long bones following fracture (Blecher et al., 2017b).

Peripheral information detected by PSNs is processed via both local and long-range neural circuits. Locally, PSNs from the muscle spindle (type Ia afferents) or the Golgi tendon organ (type Ib afferents) form monosynaptic or disynaptic connections respectively with α-motor neurons which innervate the same, or functionally similar, muscle groups from which the sensory information arose (Bikoff et al., 2016; Mears and Frank, 1997; Ozaki and Snider, 1997). In contrast to this relatively well characterised local spinal reflex circuit, the long-range circuits for higher order integration prior to motor output modulation have received less attention, particularly in mouse. Early studies in cat have shown that type Ia, Ib and type II muscle PSN afferents synapse on spinocerebellar (SC) neurons (SCNs) of the spinal cord that send axonal projections directly to the cerebellar cortex via spinocerebellar tracts (SCTs) (Aoyama et al., 1988; Edgley and Jankowska, 1988). The more recent topographic mapping of SC projections to the cerebellar cortex in mouse (Sengul et al., 2015), which is broadly consistent with extensive anatomical literature in other species (Grant and Xu, 1988; Grant et al., 1982; Matsushita and Hosoya, 1979), provides a comprehensive understanding of where in the mouse spinal cord SC neurons reside. This study defined 5 axially-restricted populations spanning cervical (Central Cervical Nucleus (CeCV)), thoracic (Dorsal Nucleus, here referred to as Clarke’s Column (CC)), lumbar (Lumbar Precerebellar Nucleus (LPrCb) and Lumbar Border Nucleus (LBPr)) and sacral (Sacral Precerebellar Nucleus (SPrCb)) regions, and 4 scattered populations spread along the rostro-caudal (R-C; top-to-bottom) extent of the spinal cord within the deep dorsal horn (DDH) and laminae VI-VIII (Sengul et al., 2015). These populations are of mixed embryonic origins, with a small subset of CC cells and various scattered populations deriving from an *Atoh1*+ lineage (Bermingham et al., 2001; Yuengert et al., 2015), others from a *Neurog1*+ lineage (Sakai et al., 2012) and the remaining majority of SCN of unknown origin. How these mixed-lineage populations are divided among the regional nuclei of the SC system is currently unknown.

The Hox family of transcription factors are fundamental regulators of development across bilateria, endowing cells with the positional information required for region-specific morphology and functionality (Wellik, 2009). In mouse and human, 39 Hox genes are subdivided into 13 paralogous groups (*Hox1*-*13*) based on evolutionary descent, with varying subsets of the 13 paralogs genomically clustered at 4 distinct loci (*HoxA-D* clusters). Transcription from each *Hox* cluster is, however, not limited to *Hox* protein-coding genes. A vast array of noncoding transcripts are also produced, including microRNAs and long-noncoding RNAs (reviewed in Casaca et al., 2018). Developmental expression of these noncoding RNAs is largely consistent with the ordered spatio-temporal activation of *Hox* gene expression that occurs from the 3’ to 5’ ends of each cluster (Mansfield and McGlinn, 2012; Mansfield et al., 2004; Wong et al., 2015). Moreover, important developmental functions have been ascribed to many of these Hox-embedded noncoding RNAs (reviewed in Casaca et al., 2018 and Heimberg and McGlinn, 2012), including a striking homeotic role for the miR-196 family of miRNAs (McGlinn et al., 2009; Wong et al., 2015).

The central importance of Hox genes in imparting positional identity is particularly apparent in the developing nervous system, where cell-intrinsic mechanisms dictating migration and connectivity can change dramatically dependent on the Hox code of a neuron (Bechara et al., 2015; Gavalas et al., 1997; Mark et al., 1993; Oury et al., 2006; Studer et al., 1996; reviewed in Philippidou and Dasen, 2013). Prominent examples of this in the spinal cord are the generation of axially-restricted motor neuron (MN) columns which harbour neurons that project to broad muscle target regions (e.g. limb versus axial musculature), and the more restricted MN pools within these columns, which target specific muscles within a region. A combinatorial Hox code correlates with the axial restriction of column and pool subtypes, and moreover, functionally defines their appropriate connectivity (Dasen et al., 2003, 2005; Philippidou and Dasen, 2013). The most striking example is seen following loss of *Hoxc9*, where the trunk-specific hypaxial motor column is completely lost and the lateral motor column, normally specific to brachial and lumbar levels, subsequently extends along the entire intervening R-C axis (Jung et al., 2010). Pool identity is also affected in this and other *Hox* mouse knockouts, with neurons displaying random organisation within a motor column, often projecting to more anterior muscle targets (Jung et al., 2010; Wu et al., 2007).

Here, we describe widespread expression of *Hox*-cluster genes throughout SC neurons of the thoracic, lumbar and sacral regions. This expression is seen from early embryonic stages through to late adulthood, indicating their potential to function in the genesis and homeostasis of SC circuits. We find that the spatial regionalisation of Hox code along the R-C axis parallels the anatomical subdivision of SC neurons within the spinal cord, and that loss of an individual Hox gene leads to a dramatic reduction of an axially-restricted SC subtype. This work suggests a commonality in spinal cord patterning mechanisms across diverse neuron types and provides critical molecular tools that allow early and comprehensive visualisation of SC development.

## Results

### Posterior *Hox*-cluster reporter expression is observed in the adult and developing cerebellum

We have previously generated a mouse knock-in reporter line that drives expression of eGFP from the endogenous *miR-196a2* locus, genomically positioned between *Hoxc9* and *Hoxc10* (referred to as *196a2*-eGFP) (Wong et al., 2015). Detailed characterisation of this reporter line revealed unexpected eGFP protein within the adult and neonatal cerebellum, far outside the posteriorly-biased pattern of expression one would expect from a gene located within the posterior *HoxC* cluster.

Analysis of *196a2-*eGFP by immunofluorescence at postnatal day (p) 28 revealed a highly reproducible and restricted pattern of expression forming rosettes within the granule-cell layer of cerebellar lobules I-V, VIII and IX (Figure 1A), which clustered in parasagittal stripes as shown in coronal section (Figure 1B; summarised in Figure 1C). The observed rosettes, lobule restriction and parasagittal banding correspond closely with published literature on the terminal projection pattern of spinocerebellar (SC) neurons in mouse (Reeber et al., 2011). One notable exception to this was reproducible eGFP expression on the ventral side of lobule IX in the most posterior region (Figure S1A), a site commonly associated with vestibular input, that we find no description of in any historical SC anterograde tracer literature. However, the lack of eGFP detection within vestibular nuclei (Figure S1B), together with results of more recent single SC axon tracing (Luo et al., 2018), support our identification of eGFP at this site as originating from a spinocerebellar projection. The absence of eGFP expression in cerebellar nuclei (Figure S1C) indicates these *196a2*-expressing neurons do not send collaterals to this location as has been observed for some SC neurons in rat (Matsushita, 1997, 1999a, 1999b; Matsushita and Xiong, 1997; Matsushita and Yaginuma, 1995) and mouse (Luo et al., 2018). This pattern of eGFP expression was maintained from neonatal through to late adult life (Figures S1A, D-F), with expression of eGFP observed in fibres of passage in two distinct tracts corresponding to the location of the ventral (VSCT) and dorsal (DSCT) SC tracts (Figure S1D), and the stereotypical parasagittal banding readily apparent at 1 week of age. At all stages, eGFP protein did not appear perinuclear within local cerebellar cells as determined by DAPI staining (data not shown), nor was *eGFP* mRNA detected by *in situ* hybridisation within the cerebellum (data not shown), supporting the notion that *196a2-eGFP*-expressing cells originate outside the cerebellum, with eGFP protein filling axonal projections and synaptic terminals. Parallel characterisation of a second reporter line, *miR-196a1-eGFP*, which drives eGFP expression from the paralogous gene locus of the *HoxB* cluster, also revealed expression indicative of adult SC neurons (Figure S2).

**Figure 1:**
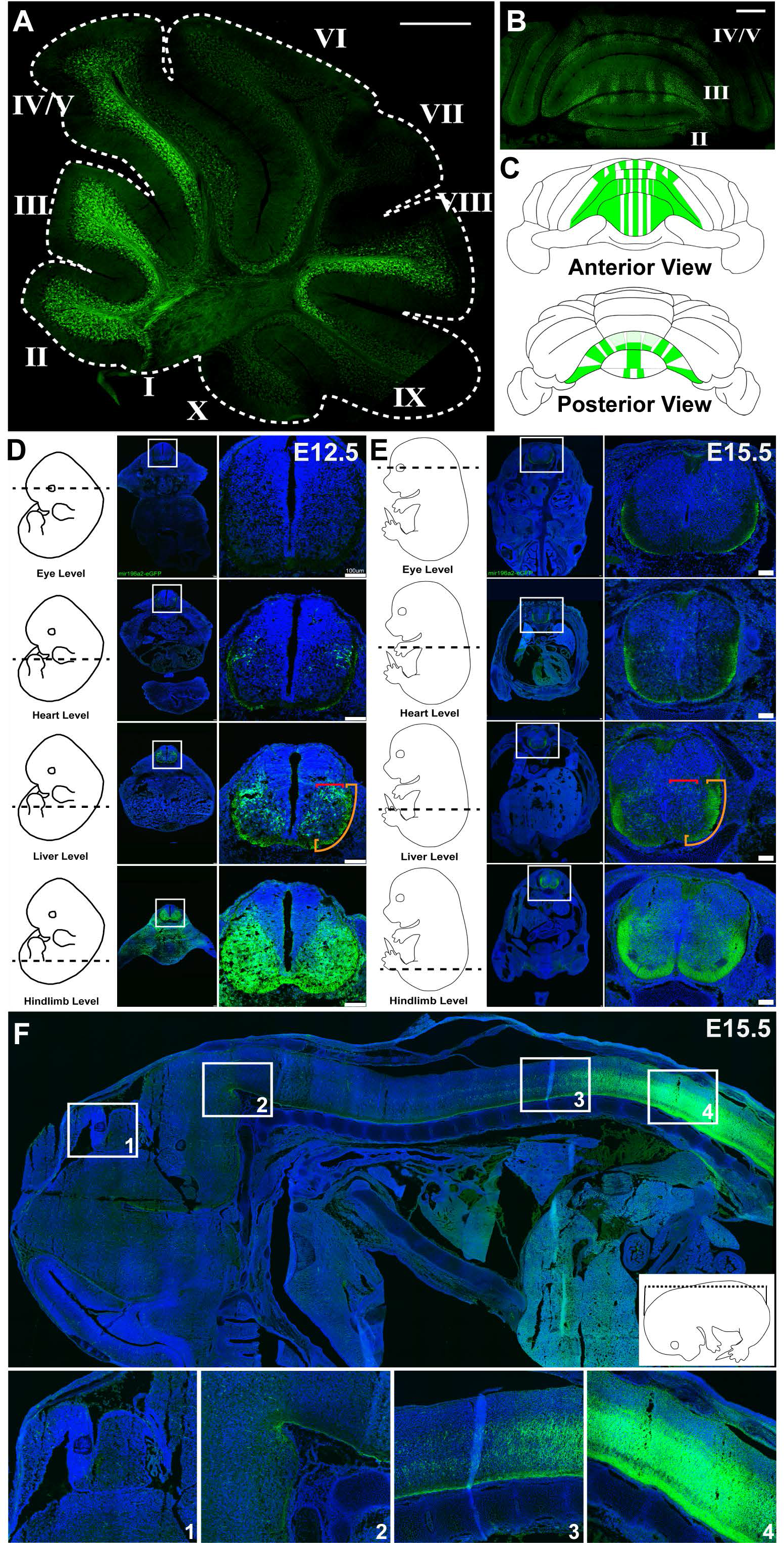
*miR196a2-eGFP* reporter expression marks putative spinocerebellar neurons. (A-F) Immunofluorescent detection of eGFP protein in adult and embryonic miR*196a2-eGFP* reporter mice. (A) Sagittal section at p28 reveals lobule-restricted eGFP in the cerebellum. (B) Coronal section at p28 reveals parasagittal banding of eGFP in the cerebellum. (C) Cartoon composite of eGFP expression along the A-P axis of the cerebellum, based on complete coronal series from three p28 animals. (D-E) Cross-sections of E12.5 (D) and E15.5 (E) embryos reveal axially-restricted eGFP expression along the A-P axis. For each series, cartoon embryos indicate the axial level where sections were imaged (dotted line) and the right panel provides a magnified view of the neural tube. Red bracket indicates the location of cell bodies and orange bracket indicates the location of SC tracts. (F) Sagittal section of an E15.5 embryo reveals eGFP-positive cell bodies in the posterior embryo and eGFP-positive neural tracts extending rostrally into the developing brain. Boxes represent regions of interest where higher magnification images in the bottom panels were taken. Scale bars (A-B, F) = 500 μm, scale bars (D-E) = 100 μm. Roman numerals I-X indicate lobules of the cerebellum.

To investigate the potential origin of cerebellar *196a2*-eGFP expression, we characterised reporter expression within the adult spinal cord. At p28, eGFP displayed region-specific expression in cell bodies along the R-C axis, first visible from approximately thoracic (T) spinal segment T3, in large scattered neurons of laminae 5-7 and small neurons in lamina 1 (Figure S3). The number of neurons in the medial region increased more posteriorly, with a clustered population becoming apparent in the region of CC (Figure S3). Fibre bundles positive for eGFP were observed in the region of the ascending DSCT and VSCT along the entire R-C extent of the spinal cord, particularly evident at upper thoracic and cervical levels where eGFP-positive cell bodies were not present (Figure S3).

Knowledge of the molecular signatures defining collective and individual SC neurons is scarce, a fact that underpins the current lack of molecular tools to visualise ascending SC axonal projections during development. To determine whether our *196a2*-eGFP reporter mouse can be used to visualise the genesis of SC circuitry, we characterised eGFP expression at embryonic stages corresponding to timepoints when SC axons begin to ascend towards (E12.5), or enter (E15.5), the cerebellar primordium. In the E12.5 neural tube, we observed eGFP expression in cell bodies and ventrally-biased fibre bundles up to the level of the forelimb/heart (Figure 1D). As development proceeded, these fibre bundles were seen at more rostral levels, and by E15.5 we could detect eGFP-positive tracts along the entire rostral spinal cord and hindbrain (Figures 1E and F) with a large majority of these fibres entering the cerebellar primordium (Figure 1F, inset 1). Together, the striking expression revealed by these reporter lines allow us to propose that *Hox*-cluster genes are expressed in SC neurons during both genesis and adult homeostasis.

### *miR-196a2-eGFP* is dynamically expressed across spinocerebellar neuronal columns

We next investigated whether *196a2*-eGFP was expressed throughout all major SC nuclei and scattered populations, or was restricted to subpopulations. To confirm that *196a2*-eGFP is in fact localised to SC neurons, we first performed dual immunofluorescent staining for eGFP alongside the vesicular glutamate transporter VGLUT2 which is strongly expressed in, though not exclusive to, SC terminals (Gebre et al., 2012). Indeed, all eGFP-expressing rosettes in the granule cell layer of p28 animals co-labeled with VGLUT2 (Figures 2A-E). Given the current paucity of molecular markers defining individual SC nuclei and scattered populations, we next performed retrograde tracing on *196a2*-eGFP postnatal pups. A glycoprotein-deleted rabies virus expressing the mCherry fluorescent reporter (*Rabies-ΔG-mCherry*; (Wickersham et al., 2007) was injected into the p2 (n=1) or p4 (n=3) *196a2*-eGFP cerebellum, and spinal cords were collected for analysis at p7 (Figure S4). Colocalisation analysis of spinal cord sections revealed mCherry/eGFP double-positive cells within almost all previously defined SC populations, including CC, LBPr and SPrCb nuclei, and scattered cells within the DDH and 8Sp (Figures 2G-J” and L-L”, see STAR methods). Exceptions to this included the CeCV nucleus which falls outside the T3-Co3 expression domain of the *196a2*-eGFP reporter and thus would not be expected to exhibit mCherry/eGFP dual-positivity. Additionally, the LPrCb nucleus lacked any evidence of mCherry/eGFP dual-positivity (Figures 2K-K” and S5). This latter observation was unexpected in light of confirmed eGFP expression in adjacent SC populations both rostral and caudal to the LPrCb nucleus (Figures 2H-H” and L-L”), and even within LBPr and scattered SC cells of the same or similar axial levels (Figure 2I-I” and J-J”). This supports the presence of unique molecular signatures across SC populations that do not rely solely on axial level, though we cannot formally exclude technical limitations given the total number of mCherry-positive LPrCb cells labelled following retrograde tracer injection was not large. Focusing on the proportion of mCherry-positive neurons which co-expressed eGFP, we observed considerable variation between the various SC populations. For example, while over 90% of mCherry-positive cells within CC and the SPrCb nucleus co-expressed eGFP, most of the scattered populations showed approximately 50% co-expression (Figure S5). Moreover, a gradient of mCherry/eGFP dual-positivity was observed along the R-C extent of CC, ranging from ~30% at upper thoracic levels (T3-T7) compared to ~95% at lower levels (T8-T13) (Figure S5). Together, these various approaches have allowed us to demonstrate expression of a *Hox*-cluster gene within SC neurons, with detailed characterisation of *196a2*-eGFP indicating heterogeneity within and between SC populations.

**Figure 2:**
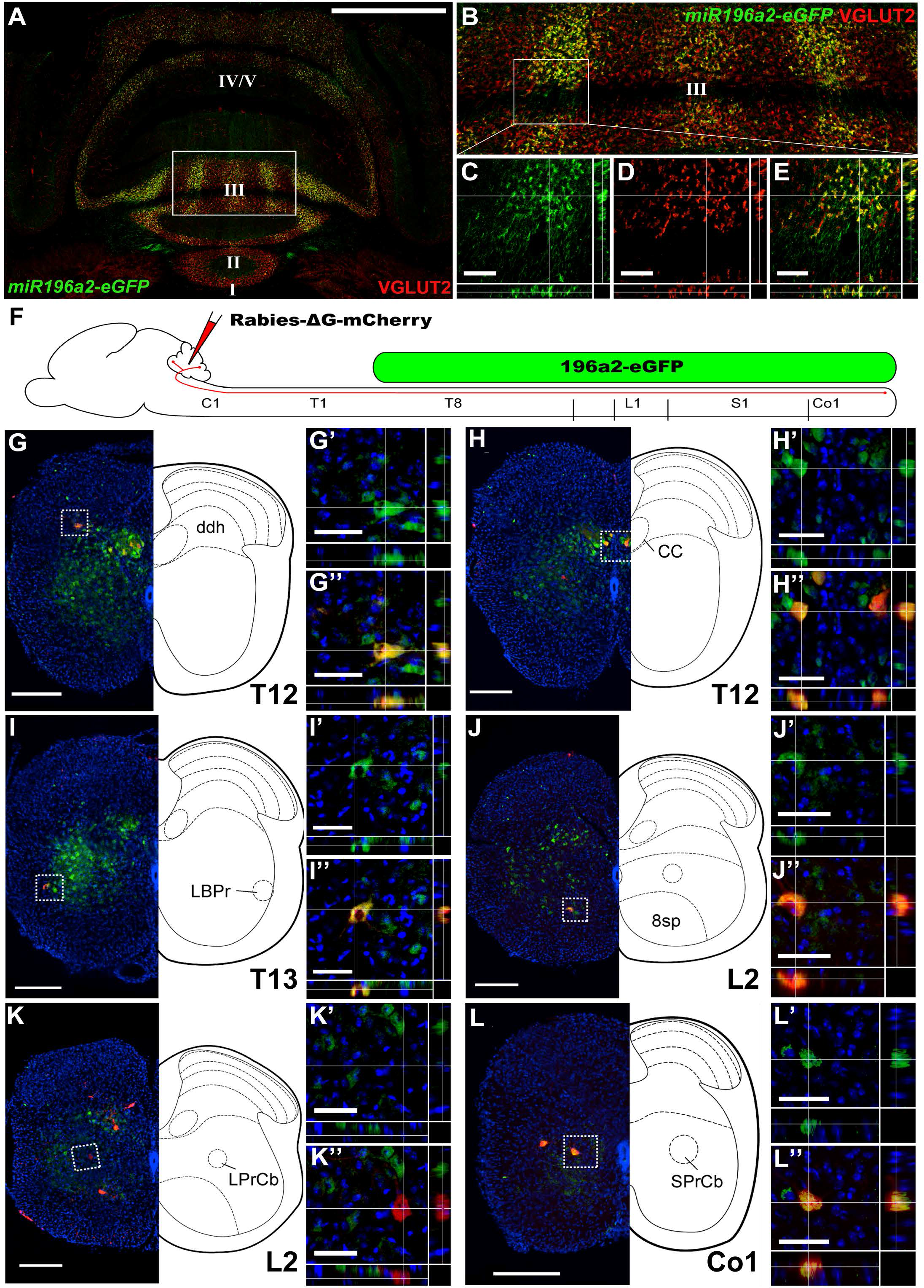
*miR196a2-eGFP* expression is confirmed within spinocerebellar neuron subpopulations. (A-E) Coronal section of a p28 *miR196a2-eGFP* cerebellum, co-stained for eGFP (green) and VGLUT2 (red) proteins. (A-B) Maximum projection images. The boxed region encompassing lobule III in (A) can be seen in higher magnification in (B). I-V = lobules of the cerebellum. Scale bar (A) = 1mm. (C-E) Z-projection view of the boxed region in (B). Visualisation of eGFP (C), VGLUT2 (D) and a merge of the two (E) demonstrates co-labelling of these proteins at synaptic terminals in 3 dimensions. Scale bar (C-E) = 50μm. (F) A schematic of the retrograde tracer experimental strategy. Approximate locations of sections used for imaging in (G-L) are marked. C, Cervical; T, Thoracic; L, Lumbar; S, Sacral; Co, Coccygeal. (G-L) Cross-section of p7 *miR196a2-eGFP* spinal cords following Rabies-◻G-mCherry retrograde tracer injection at p4. Colocalisation analysis of eGFP and mCherry was performed for each major spinocerebellar nucleus and for various scattered populations from the thoracic level and below. Cartoon diagrams were drawn from adjacent sections stained for acetylcholine esterase. White boxes indicate the location of inset Z-projection views presented for eGFP expression (G’-L’) and merged eGFP and mCherry expression (G”-L”). eGFP expression was observed in each population except the LPrCb (K-K”). Scale bars (G-L) = 200 μm, scale bars (G’-L’) and (G”-L”) = 50 μm. DDH, deep dorsal horn; LBPr, lumbar border cell; 8Sp, lamina 8 cell; CC, Clarke’s column; LPrCb, Lumbar precerebellar nucleus; SPrCb, Sacral precerebellar nucleus.

### Spinocerebellar neurons display population-specific repertoires of *Hox* gene expression

The *miR-196* genes are genomically located in intergenic regions between *Hox9* and *Hox10* paralogs (Yekta et al., 2004) and their expression conforms to the overarching vertebrate Hox-cluster constraint of collinear temporal activation during embryogenesis (Wong et al., 2015). This prompted us to question whether posterior *Hox* (*Hox9-11*) genes are also expressed in this neuronal context. To date, however, there is limited information as to whether *Hox9-11* gene expression is maintained in the developing spinal cord throughout the late embryonic and early postnatal stages critical to SC tract formation and circuit establishment. To this end, we performed a section *in situ* hybridisation screen for the *Hox9-11* genes, and found expression of all ten genes maintained in the neural tube/spinal cord from E12.5, when SC axons begin ascending rostrally, through to p7, when mature patterns of connectivity with the cerebellum have been established (Figure S6).

To assess *Hox* gene expression specifically within SC neurons, we performed an identical p7 *in situ* screen following injection of the *Rabies-ΔG-mCherry* retrograde tracer into the cerebellum of p4 wildtype (WT) pups (n=4) (Figure 3). Each *Hox9*-*11* gene was expressed in at least three SC populations, and collectively, each SC population was positive for expression of at least seven Hox genes (Figure 3C). Assessing from rostral to caudal, the CeCV nucleus is devoid of *Hox9-11*, being anterior to most of these genes’ expression domains and a location where we would predict *Hox5*-*8* paralog expression. The subsequent major SC nuclei can be characterised by distinct Hox expression profiles: CC being *Hox9*-positive in the upper thoracic region, a *Hox9/Hox10*-positive lower CC and LPrCb, and a *Hox9-11*-positive SPrCb signature. Widespread *Hox9-11* expression was seen in scattered populations, with the exception of DDH cells where no expression of *Hox10* paralogs was detected. Collectively, this analysis has allowed us to define the “Hox code” of major SC nuclei and scattered SC populations, with expression of more 5’ *Hox* genes demarcating progressively more posterior populations within the SC system.

**Figure 3:**
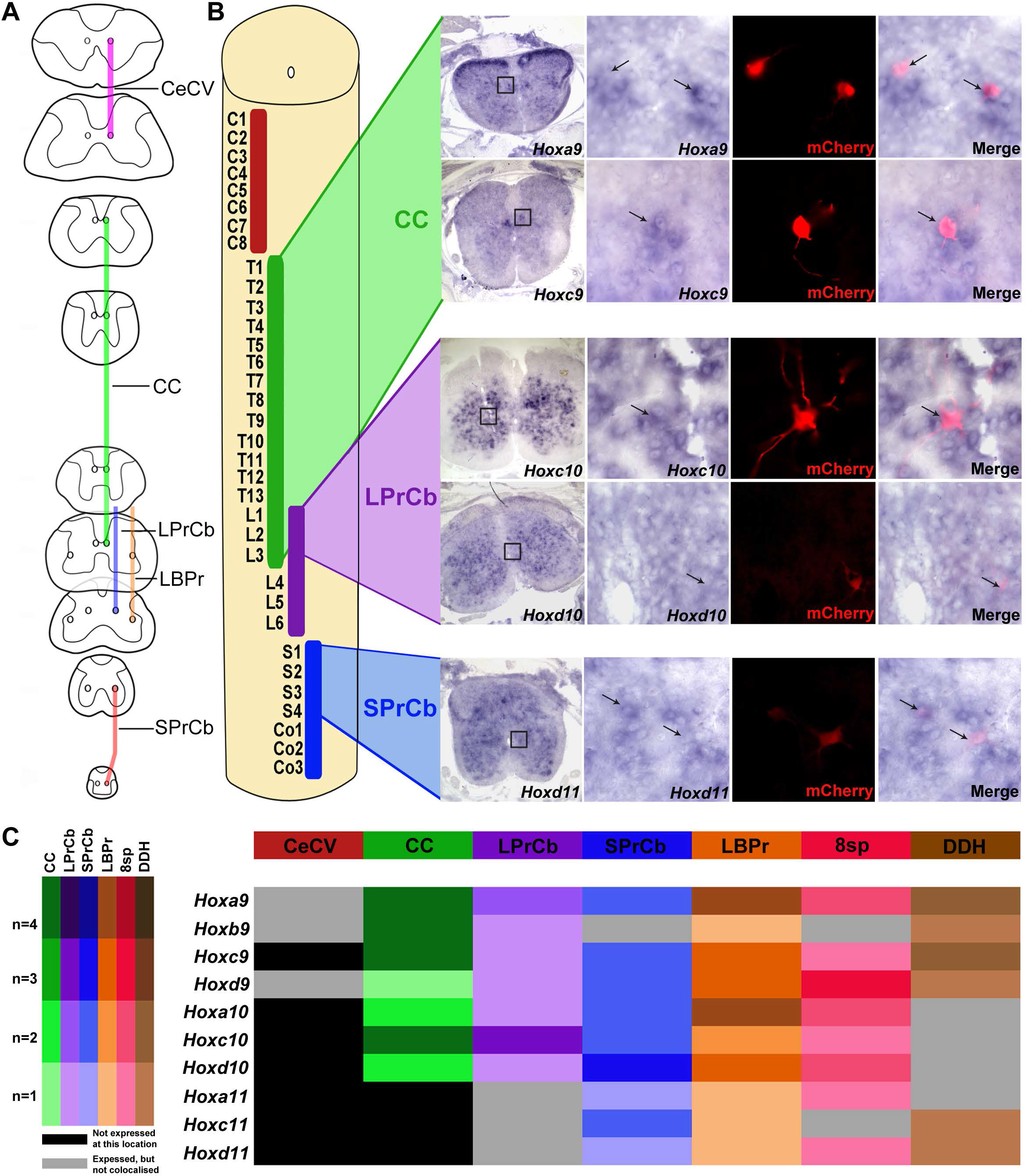
*Hox 9-11* paralogs molecularly delineate spinocerebellar subpopulations. (A) Cartoon summary of each major spinocerebellar nucleus in mouse. (B-C) An *in situ* hybridisation screen for expression of all *Hox9-11* genes in the p7 wildtype spinal cord following Rabies-◻G-mCherry retrograde tracer injection at p4. (B) The *in situ* expression pattern of selected Hox mRNAs is presented, demonstrating individual cells colabelled for a Hox mRNA and mCherry, with axial levels indicated. The location of inset images is demarcated with a black box. (C) Summary heat map cataloging the expression of each *Hox9-11* gene (rows) within each major spinocerebellar nucleus or scattered population (columns). The screen was performed on 4 animals, darker shades indicate the number of replicate animals where colabelling was observed (see legend on left). Scale bars = 200 μm (B-F) and 50 μm (B’ – F”’).

### Hoxc9 function is essential within a large subset of CC neurons

Functional analysis of Hox genes, and of Hox-embedded microRNAs such as miR-196, has been challenging due to redundancy between paralogs (McIntyre et al., 2007; Wellik and Capecchi, 2003; Wong et al., 2015). In the context of MN columnar organisation however, Hoxc9 activity alone is responsible for imparting thoracic identity and restricting limb-innervating lateral motor columns to the fore- and hind-limb levels (Dasen 2003; Jung 2010; (Baek et al., 2017)). Following detection of *Hoxc9* expression in thoracic-level SC neurons of CC (Figure 3), we proceeded to investigate the effect of *Hoxc9* removal by performing retrograde tracing experiments in *Hoxc9*^−/−^ and WT littermate mice. Fluoro-Gold (FG) tracer was stereotactically injected at 6 coordinates along the anterior-posterior extent of the cerebellum of 12 week old mice, with the aim of providing comprehensive and reproducible coverage of the vermis. Following confirmation of reproducibility between animals (Figure S8), we undertook counts of FG-labelled cells in four major SC populations: CeCV, CC, LPrCb and SPrCb; with counts for CC conducted at three axial levels corresponding to upper, mid and lower thoracic spinal cord segments (Figure 4). Comparison of *Hoxc9*^−/−^ and WT spinal cords (n = 4 animals per genotype) revealed no significant change in the number of FG-labelled neurons per section within CeCV, LPrCb or SPrCb populations (Figure 4B), consistent with their position outside of the known domain of Hoxc9 functional activity. In contrast, a significant reduction in FG-labelled neuron number within CC was observed in *Hoxc9*^−/−^ animals compared to WT at all three thoracic axial levels assessed (Figures 4A and B). Variation in the extent of tracer-positive cell reduction was observed dependent on axial level, with a maximum of ~75% reduction seen at lower thoracic regions (Figure 4B; p < 0.001). It is important to note that our results demonstrate a dramatic reduction, but not complete abolition, of FG-labelled CC cells in *Hoxc9*^−/−^ mutants. Hoxc9 function is therefore only required in a subset of CC neurons, in contrast to its requirement for specification of the entirety of thoracic-specific preganglionic and hypaxial motor columns (Jung et al., 2010).

**Figure 4:**
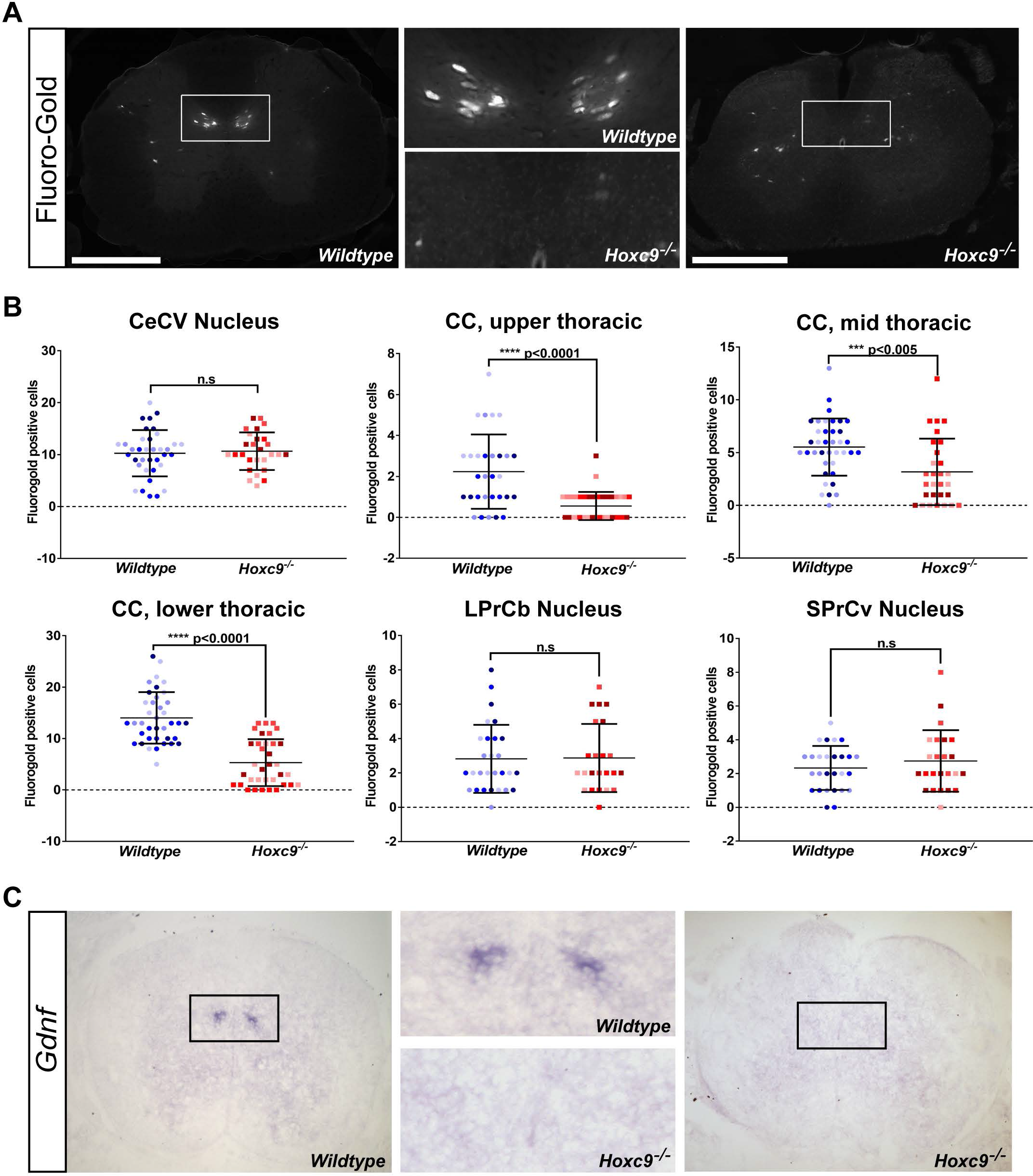
Region-specific loss of spinocerebellar neurons in *Hoxc9*^−/−^ mutant mice. (A) Tracing of cerebellar-projecting neurons following Fluoro-Gold (FG) injection into the adult cerebellum revealed a visible reduction of FG-positive cells in CC of *Hoxc9*^−/−^ animals compared to WT. Scale bars = 500 μm. (B) Quantification of FG-positive cell number per section at defined axial regions along the spinal cord showed a statistically significant reduction in FG-positive cell number across three regions of CC in *Hoxc9*^−/−^ animals compared to WT, but no change in CeCV, LPrCb or SPrCb populations (Multiple T-tests +/− SEM, two-tailed, Bonferroni-Sidak method). Data points from individual animals within a genotype are colour-coded. (C) *In situ* hybridisation for *Gdnf* in the E18.5 spinal cord revealed a complete loss of *Gdnf*-positive cells in the region of CC in *Hoxc9*^−/−^ animals compared to WT.

To corroborate retrograde tracing results, we assessed the expression of *Glial-derived neurotrophic factor* (*Gdnf*). *Gdnf* expression has been used as a marker of CC (Hantman and Jessell, 2010), though it is currently not known what proportion of CC cells express *Gdnf*, nor the upstream factor(s) driving expression within these cells. Comparison of *Hoxc9*^−/−^ and WT spinal cord sections at E18.5 (Figure 5A; n=3/3 animals per genotype) and E15.5 (Figure S9A; n=3/3 animals per genotype) revealed a complete loss of *Gdnf* in mutant embryos. *Gdnf* expression in the kidney remained unperturbed in *Hoxc9*^−/−^ embryos (Figure S9B), a site where Hox11 paralogs are known to directly regulate *Gdnf* expression (Gong et al., 2007), indicating cell type specificity and flexibility in Hox-dependent upstream regulation. Together, our work has revealed an essential role for Hoxc9 in a large majority of CC cells, and its requirement for the expression of *Gdnf* in this context.

**Figure 5:**
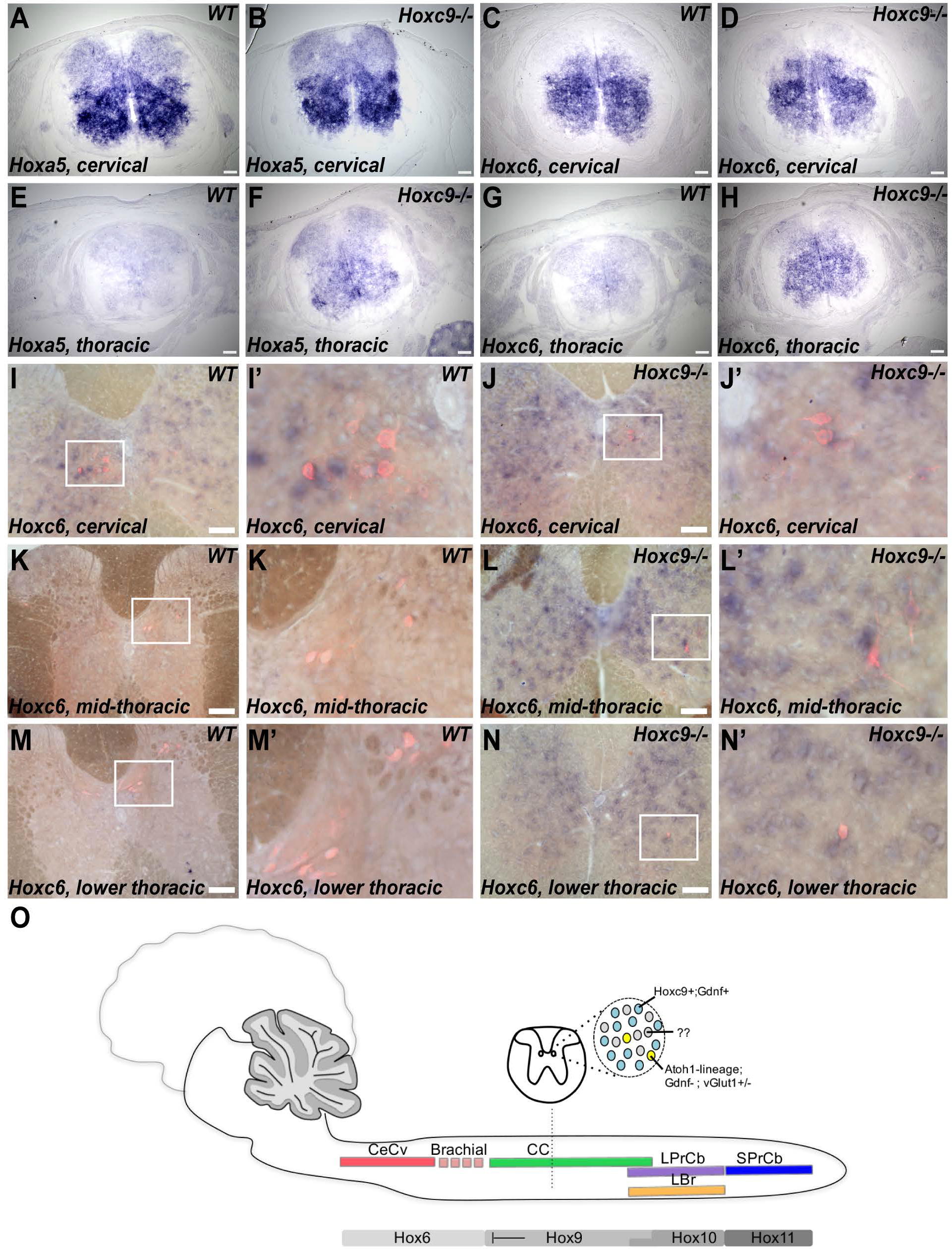
Thoracic spinocerebellar neurons display cervical *Hox* identity in *Hoxc9*^−/−^ embryos and adult mice. (A-H) *In situ* hybridisation analysis of *Hoxa5* (A-B, E-F) and *Hoxc6* (C-D, G-H) in the E15.5 neural tube reveals expansion of these genes into the thoracic region of *Hoxc9*^−/−^ embryos (B, F, D, H) when compared to WT (A, E, C, G). Scale bars = 100 μm. (I-N) Retrograde tracing using Fluoro-Gold paired with *in situ* hybridisation for *Hoxc6* on WT (I, K, M) and *Hoxc9*^−/−^ (J, L, N) adult spinal cords. At cervical levels, FG-traced cells express *Hoxc6* in both WT (I-I’) and *Hoxc9*^−/−^ (J-J’) animals. The WT thoracic spinal cord was devoid of *Hoxc6* (K-K’ and M-M’) while in *Hoxc9*^−/−^ animals, FG/*Hoxc6* double-positive cells can be seen extending into mid-(L-L’) and lower (N-N’) thoracic levels. White boxes indicate the region of higher magnification presented in adjacent panels. Scale bars = 100 μm. (O) A schematic representation of the Hox code in spinocerebellar nuclei highlighting molecular heterogeneity within Clarke’s column (CC) spinocerebellar neurons. CeCv - Central Cervical Nucleus, LPrCb - Lumbar Precerebellar Nucleus, LBr - Lumbar Border cells, SPrCb-Sacral Precerebellar Nucleus, Gdnf - Glial derived neurotrophic factor.

### Hoxc9 regulates region-specific identity of SC neurons

Concomitant with the reduction of FG-positive cells within CC cells in *Hoxc9*^−/−^ embryos, we observed ectopic FG-positive cells outside of CC at the same axial levels, particularly in laminae VI and VII (Figure 4A). The location of these cells was similar to WT retrograde tracer-positive cells immediately rostral of CC but caudal to the major CeCV nucleus that we (Figure S9E) and others have observed (Sengul et al., 2015). This suggests potential expansion of a currently ill-defined brachial level SC population, which we predict would express a brachial *Hox5*-*8* code, into more caudal levels. Hoxc9 is known to delineate the posterior boundary of brachial *Hox* expression and cell identity within ventral MNs and interneurons (Dasen et al., 2003, 2005; Sweeney et al., 2018), though the cell-type exclusivity of this regulation is currently not known. Analysis of *Hoxa5* and *Hoxa6* expression in E15.5 (Figure 5A) and E18.5 neural tube (Figure S9A) shows widespread de-repression of these genes across much of the *Hoxc9*^−/−^ thoracic spinal cord when compared to WT, suggesting multiple motor and sensory neural networks are likely to be coordinately mis-patterned in *Hoxc9* mutants. With specific focus on SC neurons, we next assessed the expression of *Hoxc6* in adult *Hoxc9*^−/−^ and WT spinal cords following FG retrograde tracer injection. At cervical levels, we observe FG/*Hoxc6* double-positive cells in both WT and *Hoxc9*^−/−^ animals in both the CeCV nucleus (Figures 5I-I” and J-J”) and in less regionalised cells of laminae VI-VII at brachial levels. *Hoxc6* expression was extinguished in WT thoracic level SC neurons (Figures 5K-K” and M-M”), while we continued to observe FG/Hoxc6 double-positive cells within laminae VI-VII of the upper and mid-thoracic regions (Figures 5L-L” and N-N”) of *Hoxc9*^−/−^ animals. It is important to note, that the increase in FG-positive (i.e. cerebellar projecting) cells observed outside of CC in *Hoxc9*^−/−^ animals does not directly correlate with the dramatic decrease in FG-positive cells within CC (Figure 4A). This suggests that while some CC cells change to a more rostral SC fate and settling position following removal of Hoxc9, a substantial proportion of Hoxc9-devoid CC cells undergo alternative consequences.

## Discussion

Proprioceptive sensory neurons provide ongoing information to the central nervous system regarding changes in muscle length, muscle tension and joint position that is essential for postural stability and controlled movement. These PSNs integrate within both local spinal reflexes and long range circuits mediated by SC neurons, though the relative contribution of each circuit to motor output modulation remains unclear. This is largely due to the limited molecular understanding of SC circuitry which has hampered identification of how axially-restricted SC populations are established in the early embryo, and has consequently limited the generation of molecular-based tools with which to interrogate the origin, development and function of this system. Here, we have demonstrated extensive and coordinated expression of *Hox*-cluster genes within SC neurons (summarised in Figure 5O) and the requirement for Hox function in establishing regional specification of SC neuronal columns.

### Coordinated *Hox*-cluster expression is observed in SC neurons

To date, there is no evidence, nor expectation, that posterior *Hox*-cluster expressing neurons are located or function within the brain. As such, our identification of *miR-196a2-eGFP* within the cerebellum was unexpected. The possibility that posterior *Hox*-cluster expressing neurons project axons directly to the brain, in some cases traversing almost the entire length of the body, has until now been overlooked. This is because visualisation of posterior *Hox* mRNAs/Hox proteins has, rightly so, concentrated on the cytoplasm/nucleus of soma at posterior axial locations. The use of a fluorescent reporter was critical in allowing us to visualise distal SC neuronal structures and define axonal projection patterns in time and space. We demonstrate expression of *miR-196a2-eGFP* in rostrally-migrating axonal tracts as early as E12.5 and can follow development of this system at all stages. *196a2-eGFP* expression in this system was constant throughout the entire course of development and adult life, and thus this resource could prove invaluable in revealing the contribution, and timing, of SC dysfunction in various genetic models of disease, such as the spinocerebellar ataxias.

We have also shown that expression within SC neurons is not limited to *miR-196a* family members, but encompasses all *Hox* genes tested to date. This SC “Hox code” parallels remarkably closely what has been observed for axially-restricted MN columns. For example, the appearance of CC coincides with that of the preganglionic and hypaxial motor columns. Whether the specification of individual SC and MN progenitors utilise shared mechanisms remains to be determined since the majority of SC progenitors are yet to be identified. Our work has revealed the maintenance of posterior *Hox*-cluster gene expression in the neural tube and spinal cord well beyond the early- to mid-gestation timepoints where Hox function is often assessed. The robust expression of both *Hox*-embedded microRNAs, and of all *Hox9-11* paralogs, until postnatal day 7 indicates their potential role in all stages of SC circuitry formation, from progenitor specification through to rostral axonal migration, circuit establishment and pruning. Moreover, the maintenance of *196a2-eGFP* (Figure S1) and *Hoxc6* (Figure 5) expression in adult SC neurons is consistent with the expression of anterior *Hox* genes in adult precerebellar nuclei of the brainstem (Hutlet et al., 2016). Together this may suggest an unappreciated role for *Hox*-cluster gene function in precerebellar network homeostasis and potentially in degenerative processes. While it is expected that the functional output of these *Hox*-cluster genes in SC neurons will be through transcriptional or post-transcriptional regulation within soma of the spinal cord, one cannot rule out the possibility that Hox-embedded miRNAs act in the distal axon and presynaptic nerve terminal (Kaplan et al., 2013), or that the homeodomain-containing proteins themselves may be secreted and act non-cell autonomously within the cerebellum (Bardine et al., 2014; Prochiantz and Di Nardo, 2015). Indeed, given the complete absence of posterior *Hox*-cluster expression within neuronal cell bodies of the cerebellum, this system may serve as an excellent model to investigate such mechanisms.

### Hox function is required to axially restrict SC populations

Here, we have used *miR-196a2-eGFP* as the first active-locus reporter of SC neurons during development and in the adult. The functional requirement for miR-196 activity within this network remains to be investigated. However, the detection of both *miR-196a* paralogs within SC neurons, and the confirmation of SC expression of *Hoxa9 and Hoxa10* genes that genomically surround the *miR-196b* locus, suggests that miR-196 double or triple knockout analysis is likely to be required to detect cellular and/or phenotypic alteration of this neural network. In contrast, we observed a marked loss of cerebellar-projecting neurons from CC, and an expansion of *Hoxc6*-positive laminae V-VII SC neurons normally restricted to brachial levels, following the removal of a single Hox gene, *Hoxc9* (Figures 3 and 5). Our data is broadly consistent with that observed for thoracic-level MNs (Dasen et al., 2003, 2005; Jung et al., 2010) and spinal interneurons (Sweeney et al., 2018), whereby Hoxc9 alone is required to repress a brachial *Hox* code, thereby delineating axially-appropriate neural connectivity. An important point of difference of our work to this well-defined model, is that the marked loss of cerebellar-projecting CC neurons in *Hoxc9*^−/−^ animals does not appear to be solely accounted for by a fate switch to a more rostral SC population. In contrast to MN columns that continuously cover the R-C extent of the spinal cord, SC neurons exhibit regional fluctuations. Spinocerebellar neurons of the upper cervical region from C1 to C5 cluster medially in the CeCV nucleus. The region between this and CC nucleus which emerges around T1/T2 is less well-defined, with scattered SC cells identified within laminae V-VII (Sengul et al., 2015). The lack of well-defined SC nucleus at this location potentially reflects a degree of divergence of ascending pathways between forelimb and hindlimb/trunk PSNs, since axons of forelimb PSNs have been shown to project directly to the external cuneate nucleus of the medulla without synapsing in the spinal cord as part of the spinocuneocerebellar tract (Abrahams and Swett, 1986; Nyberg and Blomqvist, 1982, 1984). As such, a complete fate change of CC neurons to an immediately rostral population in *Hoxc9*^−/−^ mice is more difficult to reconcile than for the MN system. The fate of “lost” CC neurons in *Hoxc9*^−/−^ mice remains to be explored, though numerous possibilities exist. A failure of progenitor specification is possible, though not strongly supported by previous Hox literature. Alternatively, these neurons may fail to differentiate from a neuronal precursor population, since Hoxc9 has been shown to induce neuronal differentiation in neuroblastoma (Mao et al., 2011; Wang et al., 2013, 2014). Hoxc9 may regulate neuronal survival, given the essential requirement for Hoxc9 in inducing SC expression of the neurotrophic growth factor *Gdnf* during late embryogenesis (Figure 4). Finally, progenitors may be diverted to an as yet unknown fate, and project axons to regions other than the cerebellum, thus precluding labelling by the retrograde tracer.

The substantial changes to CC in *Hoxc9*^−/−^ mice could reasonably be expected to result in overt locomotor or behavioural defects. In our hands, the majority of *Hoxc9*^−/−^ mice die at birth, particularly when bred onto a C57B6 background. For those *Hoxc9*^−/−^ mice that survive to adulthood, many display a hunched back (Suemori et al., 1995) and our own observations) potentially representing peripubertal scoliosis, which has been identified following loss of Er81-positive PSNs (Blecher et al., 2017a), though alternative explanations such as loss of hypaxial motor column (Jung et al., 2010) may contribute to or underlie this phenotype. Similarly, the analysis of additional *Hox* gene contributions to SC function may, in many cases, be confounded by MN phenotypes, such as the hindlimb paralysis observed in *Hoxc10/d10* mutant animals (Wu et al., 2007).

### Molecular heterogeneity of SC circuitry

Our work demonstrates divergent Hox signatures that can discriminate axially-restricted SC populations (summarised in Figure 5O). Moreover, we observe molecular heterogeneity across the R-C extent of CC consistent with colinear *Hox*-cluster activation: a Hoxc9-positive upper region, an increasing proportion of *mir-196a2*-positive cells caudally, and Hoxc10- positive cells restricted to the lower regions. As Clarke’s column neurons are heterogeneous in their inputs (Hantman and Jessell, 2010; Shrestha et al., 2012), it is possible that this heterogeneity in gene expression defines neurons of specific modality, for example, *196a2*-eGFP expression in CC cells that receive hindlimb but not trunk proprioceptive input.

The extensive anatomical literature relating to SC neurons is based largely on tracer technology and lacks the precision of modern molecular tools. As a consequence, there is limited understanding of how SC circuitry is established in the developing embryo. While one progenitor population has been identified (Yuengert et al., 2015), the diversity of progenitor populations within and between SC populations is not known. Our work indicates that a large portion of CC cells are derived from a Hoxc9-positive progenitor, that during late embryogenesis require Hoxc9 to activate expression of *Gdnf*. In parallel, a separate Atoh-1 descendent population constitutes a further 10% of CC, with 0.5% of these positive for vGlut1 (Yuengert et al., 2015). Atoh-1 function is not required for the expression of *Gdnf* (Yuengert et al., 2015), further delineating these two populations (CC heterogeneity summarised in Figure 5O). Whether Atoh-1 descendents constitute all remaining CC cells in *Hoxc9*^−/−^ mice (Figure 4), or whether there are additional, molecularly-diverse, CC subpopulations, remains to be determined.

### Topography of SC projections

A great deal of research on the cerebellum has been focused on its complex molecular parcellation, and the association between this organisation and convergence of inputs from several CNS regions and cell-types based on their function (Apps and Hawkes, 2009; Hesslow, 1994; Odeh et al., 2005). Mouse mutants of Engrailed1 and Engrailed2 show widespread disruption of molecular organisation in the Purkinje cells of the cerebellum (Sillitoe et al., 2008), with corresponding alterations in the projection pattern of SC neurons to these regions (Sillitoe et al., 2010). However, the molecular networks existing within SC neurons themselves, that allow appropriate connectivity with the diverse pool of Purkinje cells, and thus formation of a topographic map within the cerebellar cortex, are yet to be determined. Similarly, there is evidence for topographic positioning of ascending SC tracts in the spinal white matter dependent on the axial level of input (Xu and Grant, 2005), though again with limited molecular understanding. The presence of an axial-specific Hox code in SC neurons could thus provide a parsimonious explanation for the control of topographic regionalisation of these neurons both in their ascending tracts and their projection to appropriate cerebellar target regions, and provide a means with which to dissect these key questions in future.

## Supporting information

Coughlan_Garside_SI

## Acknowledgements

We are grateful to George Paxinos and members of the Bourne Lab for generously providing resources and advice. We thank Charles Watson for advice and comments on the manuscript. We thank Cliff Tabin, Marian Ros, Tim Thomas and Tom Jessell for provision of plasmids for *in situ* hybridisation, Rob Bryson-Richardson for writing an ImageJ script for image correction and Deneen Wellik for providing *Hoxc9*^−/−^ mice. We thank Monash Micro- Imaging, Monash Animal Research Platform and the Monash Gene Modification Platform for technical support. This work was supported by Australian National Health and Medical Research Council Project Grant APP1063967 to E.M. F.M.R. was supported by the Swiss National Science Foundation (31003A_149573 and 31003A_175776) and the Novartis Research Foundation. The Australian Regenerative Medicine Institute is supported by grants from the State Government of Victoria and the Australian Government.

## Author Contributions

Conceptualisation, methodology and formal analysis E.C., V.G. and E.M.; Investigation, E.C., V.G., S.F.L.W., H.L., D.K., K.K., U.M. and E.M.,; Writing – Original Draft, E.C. and E.M.; Writing – Review & Editing, all authors; Funding Acquisition, E.M.; Supervision, E.M., J.B and F.R.

## STAR Methods

### Animal experimentation ethical approvals

All animal procedures were performed in accordance with the Australian Code of Practice for the Care and Use of Animals for Scientific Purposes (2013) or approved by the Veterinary Department of the Canton of Basel-Stadt. Experiments were approved by the Monash Animal Ethics Committee under project numbers MARP/2011/012 and MARP/2012/180, and the University of NSW Ethics Committee under project number 14/94A.

### Mouse strains

The following previously published mouse lines were used in this study: *miR-196a1-GFP* and *miR-196a2-GFP* (Wong et al., 2015) maintained on a C57B6 background and *Hoxc9*^−/−^ (McIntyre et al., 2007) maintained on a mixed background.

### Retrograde tracing

Neonatal injection of *Rabies-ΔG-mCherry* (Wickersham et al., 2007) into WT or *miR-196a2-GFP* animals was conducted as previously described (Di Meglio et al., 2013). Briefly, for injections in the cerebellum, p2-p4 pups were anaesthetized using isoflurane. Virus was injected using a microinjector (Narishige, IM-9B) at multiple places within cerebellar vermis to label cerebellar-projecting neurons. Less than 0.5 μl of virus (2×10^6^-10^8^ transducing units/μl) was injected per site. After injection, pups were recovered on a hot plate and then transferred back to mother. At p7, pups were perfused.

For adult injections (WT and *Hoxc9^−/−^*), Fluoro-Gold was injected at 6 locations along the rostrocaudal extent of the cerebellum per animal. Injections were performed 0.5mm lateral to the midline, at 0.5mm intervals between −5.5mm and −8.0mm caudal to the bregma, At each location, 20-40 nL of Fluoro-Gold was injected at the ventral part of the cerebellum (2-2.5mm deep from the dorsal surface), and then at 0.5mm intervals up to the dorsal surface. After the last injection, the needle was maintained *in situ* for 10 minutes before being slowly withdrawn. At the end of the procedure, skin was sutured, buprenorphine injected subcutaneously, and topical tetracycline applied over the incision site.

### Animal perfusion

Mice were anaesthetised by intraperitoneal injection of 5-20 mg sodium pentobarbital (Lethabarb, Lyppard Australia). Following confirmed absence of a toe-pinch reflex, animals were perfused intracardially with PBS at 37°C, followed by cold 4% PFA/PBS. 5 mL of each solution was used for perfusion of neonates, and 10-20 mL for adult animals.

### Dissection and tissue processing

Following perfusions, brains and spinal columns were dissected whole and fixed overnight in 4% PFA/PBS at 4°C. Samples were washed twice for 10 minutes in 0.1M PBS at 4°C, after which spinal cords were dissected from the enclosing vertebrae (for adult) and segmented. Samples were equilibrated through a series of sucrose solutions (5%, 20%, 30% sucrose in PBS) before being embedded in OCT medium (TissueTek), frozen in a bath of dry ice + methylbutane, and stored at −80°C. Frozen tissue was sectioned at 20 μm or 40 μm thickness on a Leica cryotome. Sections were either mounted directly onto slides (Superfrost, Fisher Scientific) or collected as floating sections in 24- or 48- well plates (Corning), stored in cryoprotectant (30% ethylene glycol, 20% glycerol, 50% 0.1M PBS) at −20°C.

### Histological staining

Nissl and Acetylcholine-esterase staining protocols were performed on slide-mounted sections as previously described (Homman-Ludiye et al., 2018).

### Immunofluorescence

Floating sections were rinsed in PBS, then placed in blocking solution I (5% heat inactivated goat serum, 0.5% bovine serum albumin, 0.2% Triton-X in 0.1M PBS) for a minimum of 30 minutes. Sections were incubated overnight at 4°C in primary antibody diluted in blocking solution I at specified concentrations. A full list of antibodies used is provided in the Supplementary Experimental Procedures. Following overnight incubation, sections were washed twice in blocking solution II (as above, with Triton-X replaced by Tween), then incubated with secondary antibody diluted in blocking solution II at 1:1000 concentration for 1-2 hours at room temperature. Sections were then washed twice in PBS + 0.2% Triton-X (PBTx), stained for 2-5 minutes in DAPI (1:1000 in 0.2% PBTx) and mounted on slides with ProLong Gold (Thermo Fisher) mounting media.

Slide-mounted sections were blocked for at least 30 minutes in blocking solution (as above, with 0.1% Triton-X), then incubated overnight at 4°C, coverslipped with primary antibody at specified concentration. Following incubation, slides were rinsed in PBTx (0.1%) to remove coverslips, then washed twice in blocking solution, and incubated with secondary antibody in blocking solution (1:1000) for 1-2 hours at room temperature. Slides were washed twice in PBTx (0.1%), then stained with DAPI (1:1000 in PBTx), rinsed in water, and mounted with ProLong Gold mounting media.

Immunofluorescence following *in situ* hybridisation was performed identically to standard slide-mounted procedures described here, except that PBTx was replaced in all washes by MABT.

### Section *in situ* hybridisation

Section *in situ* hybridisation was performed as previously described (McGlinn et al., 2019).

### Imaging and image processing

Wide-field brightfield images, including Nissl and section *in situ* hybridisation images, were acquired using an Olympus DotSlide with BX51 microscope and motorised stage, at 10x magnification. Other brightfield and fluorescent images were acquired using a Zeiss AxioImager Z1 with AxioCam HRm camera, and merged using Adobe Photoshop CS4. Confocal images were acquired using a Nikon C1 (Upright or Inverted) and Leica SP5 microscope, with 20x multiple-immersion or 60x oil objectives. For wide-field confocal images, 20x images were obtained in 5 Z-planes at 5μm intervals, saved as individual tiff files for each X/Y/Z coordinate, uniformly processed to remove uneven-illumination artefacts, merged into maximum projection images, and stitched using the in-built Fiji plug-in (Preibisch et al., 2009). Tiled images were acquired at 20x and stitched using Leica software.

### Cell counts and statistical analysis

Cell counts for SC nuclei are an average of 10 sections per animal per genotype for each of the thoracic regions (CC upper, mid, lower) and an average of at least 4 sections per animal per genotype for other major nuclei including CeCV, LPrCb and SPrCb. Data sets for comparison of WT and *Hoxc9*^−/−^ mutants analysed for significance using Multiple t-tests, two-tailed, Bonferroni-Sidak Method.

