## Supplementary material for "A Hox code defines spinocerebellar neuron subtype regionalisation": Coughlan_Garside_SI

**Supplemental Figure legends**

**Figure S1: Detailed characterisation of *miR-196a2-eGFP* in the postnatal cerebellum.**

(A) Confocal images of 40 µm sagittal sections of the *miR-196a2-eGFP* cerebellum at p2, p7, p28 and p68 identifies eGFP expression on the ventral side of lobule IX in the most posterior region.

(B) The observed ventral-posterior lobule IX *miR-196a2-eGFP* expression does not originate from vestibular nuclei, which show no evidence of eGFP-positive cells. Adjacent section stained by Nissl method to delineate nuclei. SPIV/LAV/MV = Spinal, Lateral and Medial Vestibular Nuclei; icp = inferior cerebellar peduncle.

(C) No eGFP-positive terminals are observed in cerebellar nuclei. Adjacent section stained by Nissl method to delineate nuclei. DN/FN/IN = Dentate, Fastigial and Interposed Nuclei.

(D) eGFP-positive fibres projecting to the cerebellum are visible in both ventral (VSCT) and dorsal (DSCT) spinocerebellar tracts from as early as p2. Nuclei stained with DAPI. Inset cartoon depicts the approximate position of section in the coronal plane.

(E-G) eGFP expression is detected in the *miR-196a2-eGFP* cerebellum at p2 (E), p7 (F) and p68 (G). I-X = lobules of the cerebellum.

Scale bars = 500 µm (B-G) and 50 µm (A).

**Figure S2: Detailed characterisation of *miR-196a1-eGFP* in the postnatal cerebellum.**

(A-E) Confocal images of 40µm sections of the cerebellum, following immunofluorescent staining for eGFP protein in *miR-196a1-eGFP* homozygous animals. Sagittal section revealed eGFP is barely detectable at p2 (A), faintly detected at p7 (B), and clearly visible in a lobule-restricted pattern at p28, consistent with SC neuron projection pattern. Coronal section at p28 revealed parasagittal banding of eGFP in the cerebellum (D-E). Insets in (D) and (E) show approximate location of coronal sections in the sagittal plane.

(F) Cartoon composite of eGFP expression along the A-P axis of the cerebellum, based on complete coronal series from three p28 animals.

I-X = lobules of the cerebellum. Scale bars = 500µm.

**Figure S3: Characterisation of *miR196a1-eGFP* and *miR196a2-eGFP* in the p28 spinal cord.**

(A-B) Confocal images following immunofluorescent staining for eGFP protein in *miR-196a1-eGFP* and *miR-196a2-eGFP* p28 animals. Representative transverse sections from across the spinal cord are shown, with the corresponding region marked in red on the schematic cartoon. Nuclei are stained with DAPI to provide anatomical reference.

(A) *miR196a1-eGFP* expression is primarily detected in the dorsal horns, beginning at C6 and continuing to the most caudal coccygeal segments. From around T4, the expression domain expands to include scattered cells in the ventral half of the spinal cord.

(B) *miR196a2-eGFP* expression can be observed in cell bodies from the T3 level, with the number of eGFP-positive cells increasing through to lumbar levels, and continuing to the most caudal coccygeal segments. Above T3, eGFP is visible in ascending fibres in the region of the dorsal and ventral spinocerebellar tracts. Expression can be observed in laminae 5-8, as well as a population in laminae 1-2 in the superficial dorsal horn from ~T8. A population of large, bright cells is visible in the region of Clarke's Column from T8 – Lumbar levels.

Cervical, C; Thoracic, T; Lumbar, L; Sacral=S. Number indicates segment number of each region (corresponding approximately to spinal nerve).

Scale bars = 200 µm.

**Figure S4: Schematic of *Rabies-ΔG-mCherry* retrograde tracer injection sites and location of labelled neurons in *miR196a2-eGFP* spinal cords.**

Four animals were used for analysis. Tracer injection sites (top row) were mapped from a coronal series of the cerebellum. The position of all fluorescently-labelled cells within the spinal cord was mapped onto spinal cord diagrams drawn from adjacent sections stained for acetylcholine esterase. The labels below each diagram indicate the location of the sections from which the neurons depicted in the diagram have been transposed. Circles delineate where neurons were counted as part of specific nuclei. Clarke's Column, CC; Lumbar Precerebellar nucleus, LPrCb; Lumbar Border cells, LBPr; Sacral Precerebellar nucleus, SPrCb; Thoracic, T; Lumbar, L; Sacral, S; Coccygeal, Co. Red dot = mCherry-positive cell; Blue dot = mCherry/eGFP double-positive cell.

**Figure S5: Heterogeneity in *miR196a2-eGFP* expression is observed within and across spinocerebellar subpopulations.**

Variation is observed in the number of eGFP-positive cells, presented as a proportion of the total number of retrograde-labelled cells, across major SC nuclei and scattered populations. The mean of 3-4 animals is presented +/- SEM. No *miR196a2-eGFP* expression was detected in CeCV nucleus. Clarke's Column is split into upper (T1-T7) and lower (T8-L3) portions to highlight the variation observed at different axial levels. The total number of cells counted for each population: CeCv=17, CC (T1 to T7)=37, CC (T8-L3)=168, LPrCb=16, SPrCb=17, Border cell=18 and 8Sp=17.

Central Cervical nucleus, CeCV; Clarke's Column, CC; Lumbar Precerebellar nucleus, LPrCb; Sacral Precerebellar nucleus, SPrCb; Lamina 8, 8Sp.

**Figure S6: *Hox9-11* paralogs are expressed in an axially-restricted manner from E12.5 through to postnatal day 7.**

Section *in situ* hybridisation was performed to characterise expression of all *Hox9-11* paralogs at embryonic days E12.5 (A) and E15.5 (B) and postnatal days p2 (C) and p7 (D). This screen revealed axially-restricted expression of all ten *Hox* genes, maintained at all timepoints.

**Figure S7: Schematic of tracer injections used to localise *Hox* expression within spinocerebellar neurons.**

Three animals were used for analysis. Schematic of the injection site/approximate spread of tracer for each animal are shown in sagittal and coronal planes for anterior and posterior regions of the p7 cerebellum.

**Figure S8: Sites of tracer injections used to analyse spinocerebellar populations in wildtype and *Hoxc9*$^{-/-}$ animals.**

Eight animals (4 per genotype) were used for analysis. Representative cerebellar sections of the 8 animals is presented, showing the extent of Fluoro-Gold diffusion for each sample and the degree of reproducibility between injections. Fluoro-Gold fluorescence was acquired by 405 nm excitation and displayed in grayscale as white staining. Left panel cartoon indicates the approximate region from where sections presented in each row have been sampled.

Scale bars = 1 mm.

**Figure S9: *Gdnf* expression is lost and *Hoxa5* and *Hoxc6* expression expanded in the thoracic spinal cord of *Hoxc9*$^{-/-}$ mutants.**

(A-B) *In situ* hybridisation for *Gdnf* in E18.5 cross sections of the thoracic spinal cord (A) and kidney (B). Despite complete loss of *Gdnf* expression in the *Hoxc9*$^{-/-}$ spinal cord, *Gdnf* expression is maintained in the kidney.

(C-D) *In situ* hybridisation for *Hoxa5* (C) and *Hoxc6* (D) in E18.5 cross sections of the thoracic spinal cord. Increased expression of both *Hox* genes is observed in *Hoxc9*$^{-/-}$ compared to WT.

(E-F) Retrograde tracing using Fluoro-Gold paired with *in situ* hybridisation for *Hoxc6* on WT adult spinal cords at cervical levels illustrating FG/*Hoxc6* double-positive cells in the less regionalised cells of laminae VI-VII at brachial levels.

Scale bars = 100µm.

### Figure S1

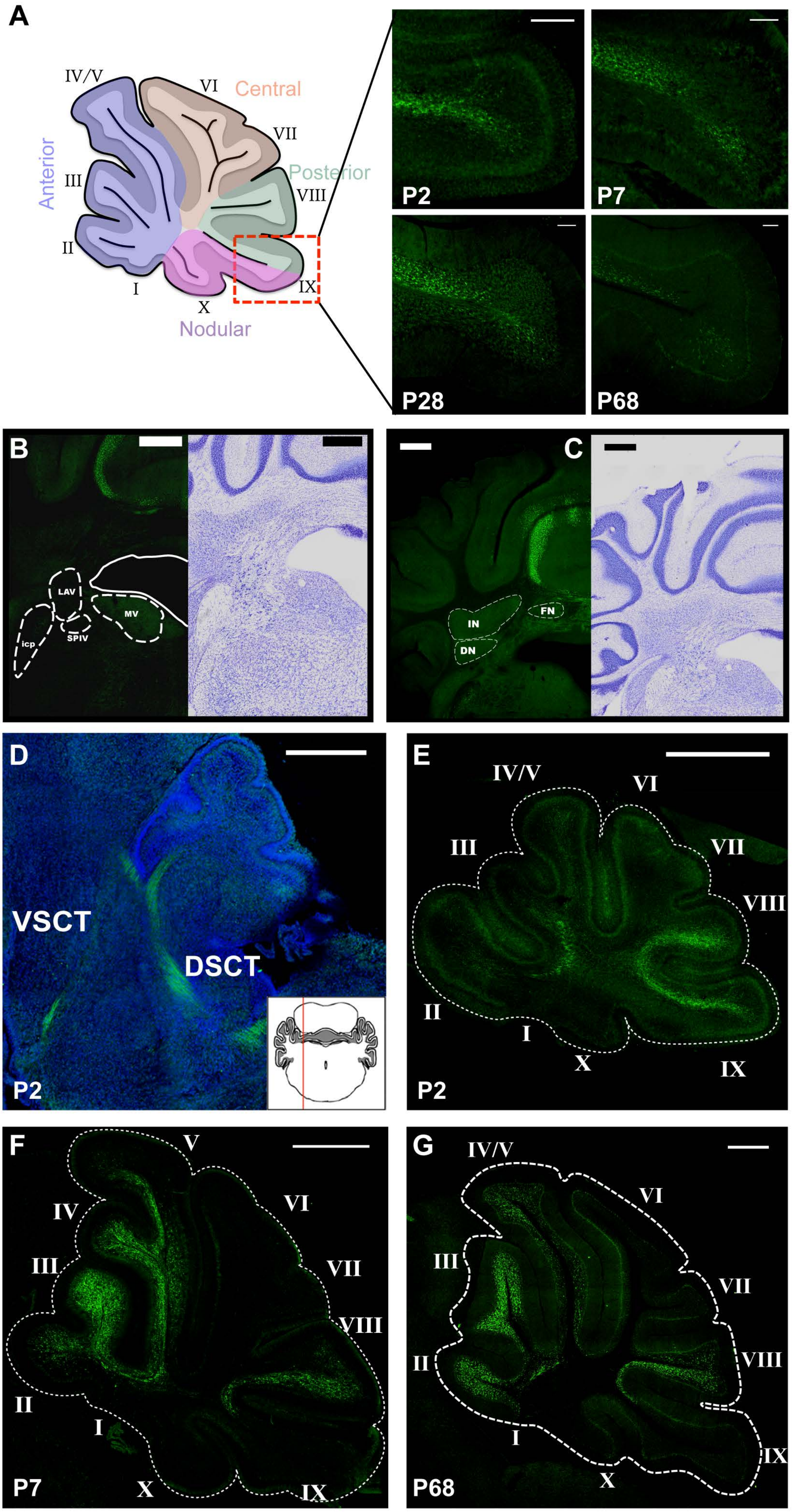
A
IV/V
VI
Central
VII
Anterior
III
Posterior
VIII
II
IX
I
X
Nodular
P2
P7
P28
P68
B
LAV
MV
icp
SPIV
C
IN
FN
DN
D
VSCT
DSCT
P2
E
IV/V
VI
III
VII
VIII
II
I
X
IX
P2
F
V
IV
VI
III
VII
VIII
II
I
P7
X
IX
G
IV/V
VI
III
VII
II
VIII
I
IX
X
P68

### Figure S2

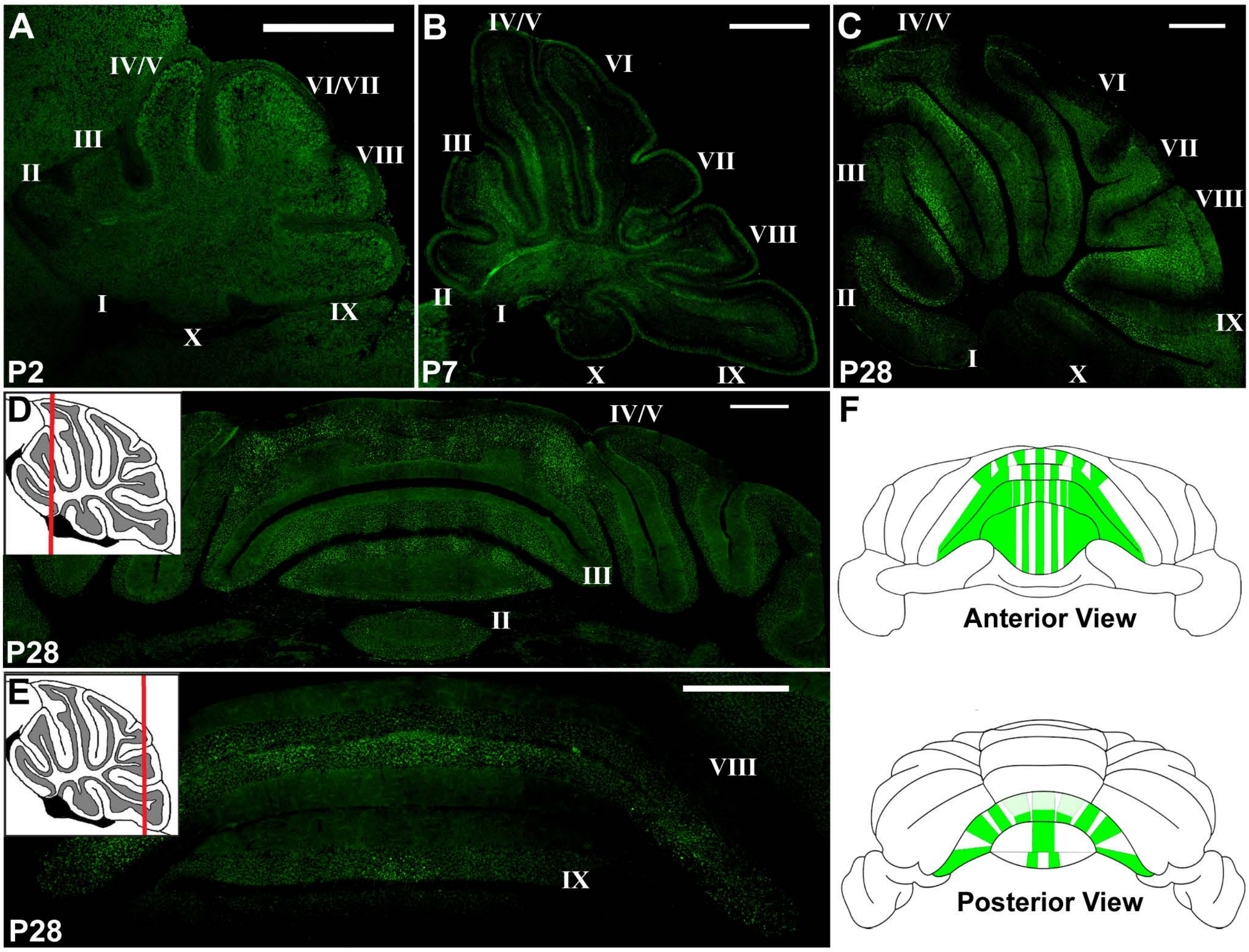

A
IV/V
VI/VII
III
VIII
II
I
IX
X
P2
B
IV/V
VI
III
VII
VIII
II
I
X
IX
P7
C
IV/V
VI
VII
III
VIII
II
IX
I
X
P28
D
IV/V
III
II
P28
E
VIII
IX
P28
F
Anterior View
Posterior View

### Figure S3

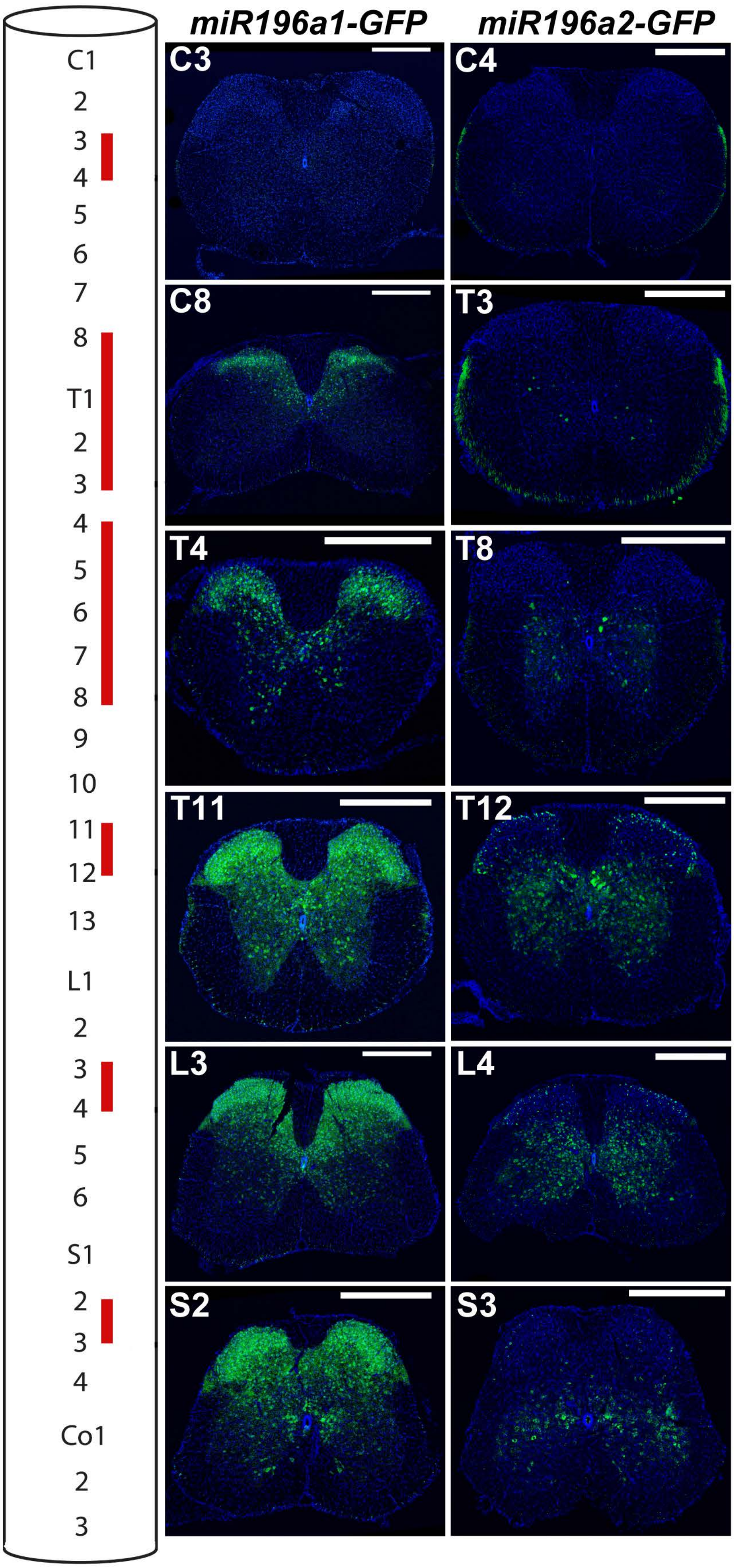
miR196a1-GFP
miR196a2-GFP
C1
2
3
4
5
6
7
8
T1
L1
2
3
4
5
6
S1
2
3
4
Co1
2
3
C3
C4
C8
T3
T4
T8
T11
T12
L3
L4
S2
S3

### Figure S4

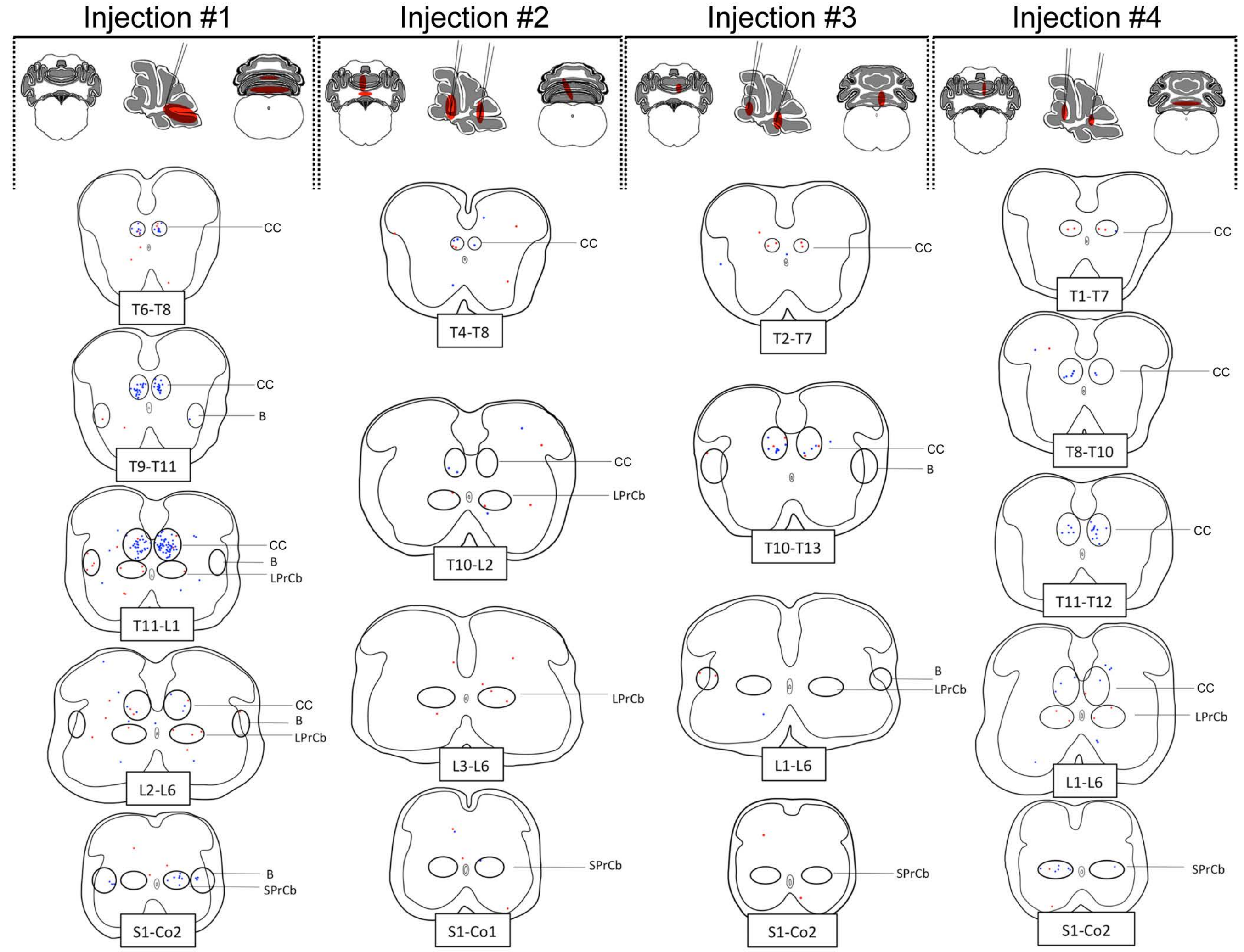
Injection #1
Injection #2
Injection #3
Injection #4
CC
T6-T8
CC
B
T9-T11
CC
B
LPrCb
T11-L1
CC
B
LPrCb
L2-L6
B
SPrCb
S1-Co2
CC
T4-T8
CC
LPrCb
T10-L2
LPrCb
L3-L6
SPrCb
S1-Co1
CC
T2-T7
CC
B
T10-T13
B
LPrCb
L1-L6
SPrCb
S1-Co2
CC
T1-T7
CC
T8-T10
CC
T11-T12
CC
LPrCb
L1-L6
SPrCb
S1-Co2

### Figure S5

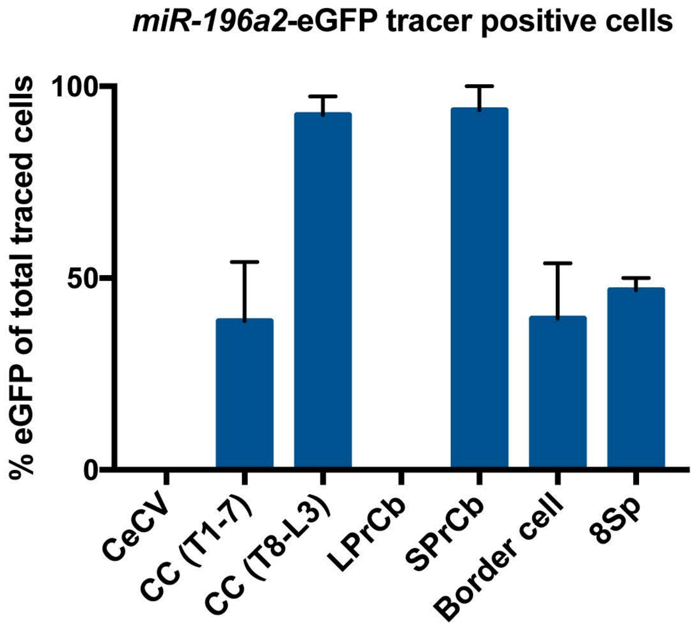
*miR-196a2*-eGFP tracer positive cells
% eGFP of total traced cells
100
50
0
CeCV
CC (T1-7)
CC (T8-L3)
LPrCb
SPrCb
Border cell
8Sp

### Figure S6

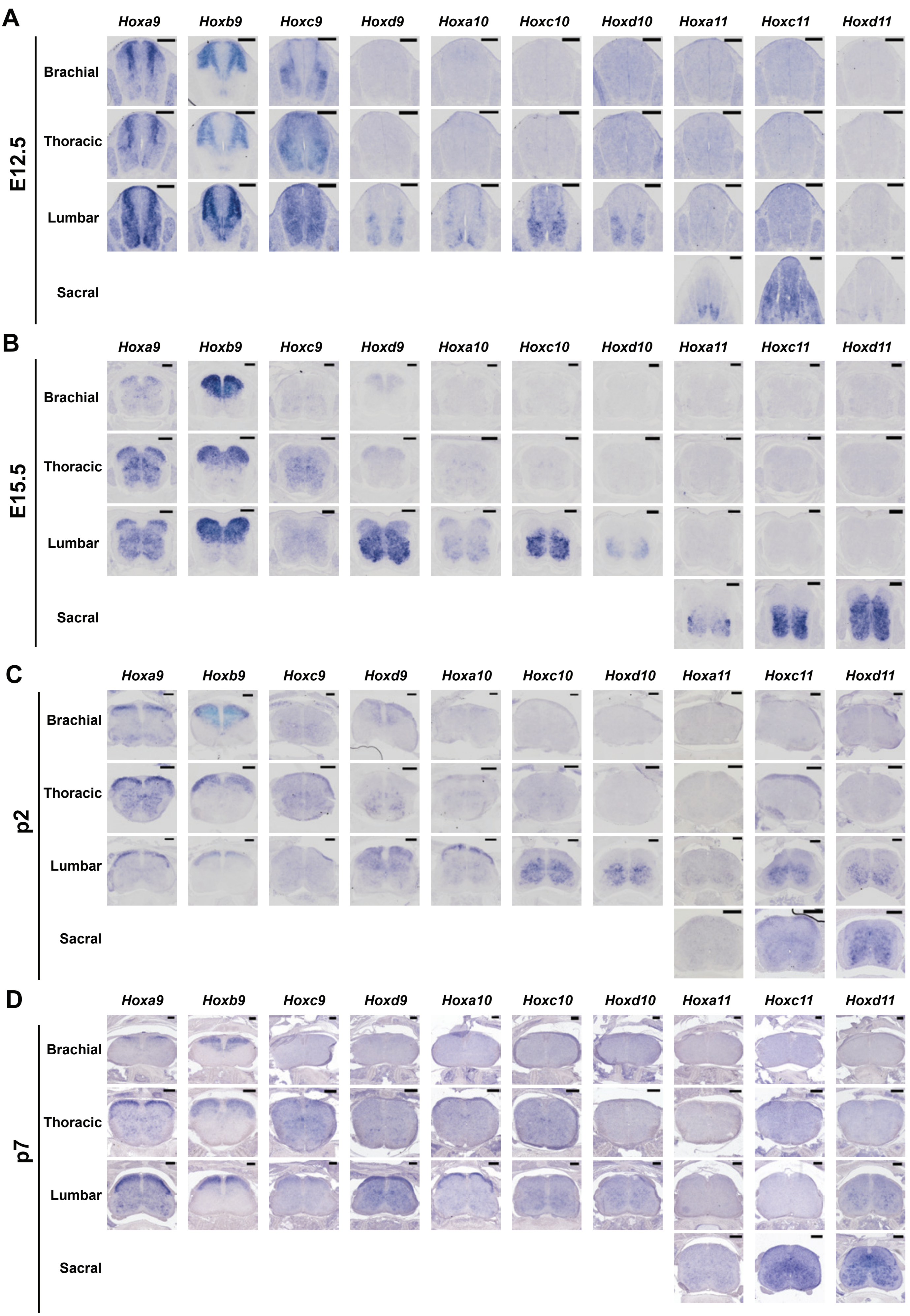
A
E12.5
B
E15.5
C
p2
D
p7
Brachial
Thoracic
Lumbar
Sacral
Hoxa9
Hoxb9
Hoxc9
Hoxd9
Hoxa10
Hoxc10
Hoxd10
Hoxa11
Hoxc11
Hoxd11

Figure S7

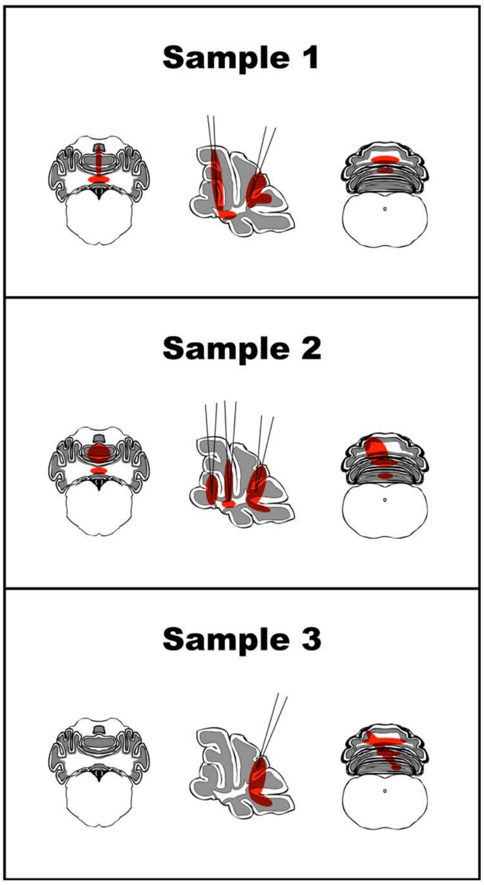

### Figure S8

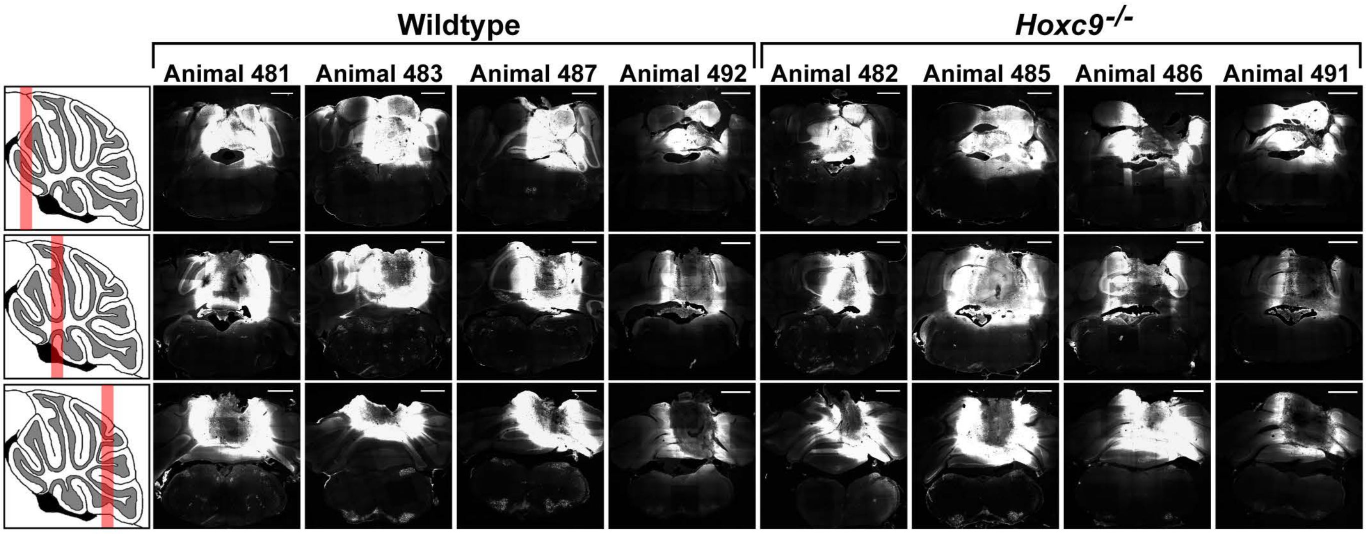
Wildtype
Hoxc9-/-
Animal 481
Animal 483
Animal 487
Animal 492
Animal 482
Animal 485
Animal 486
Animal 491

### Figure S9

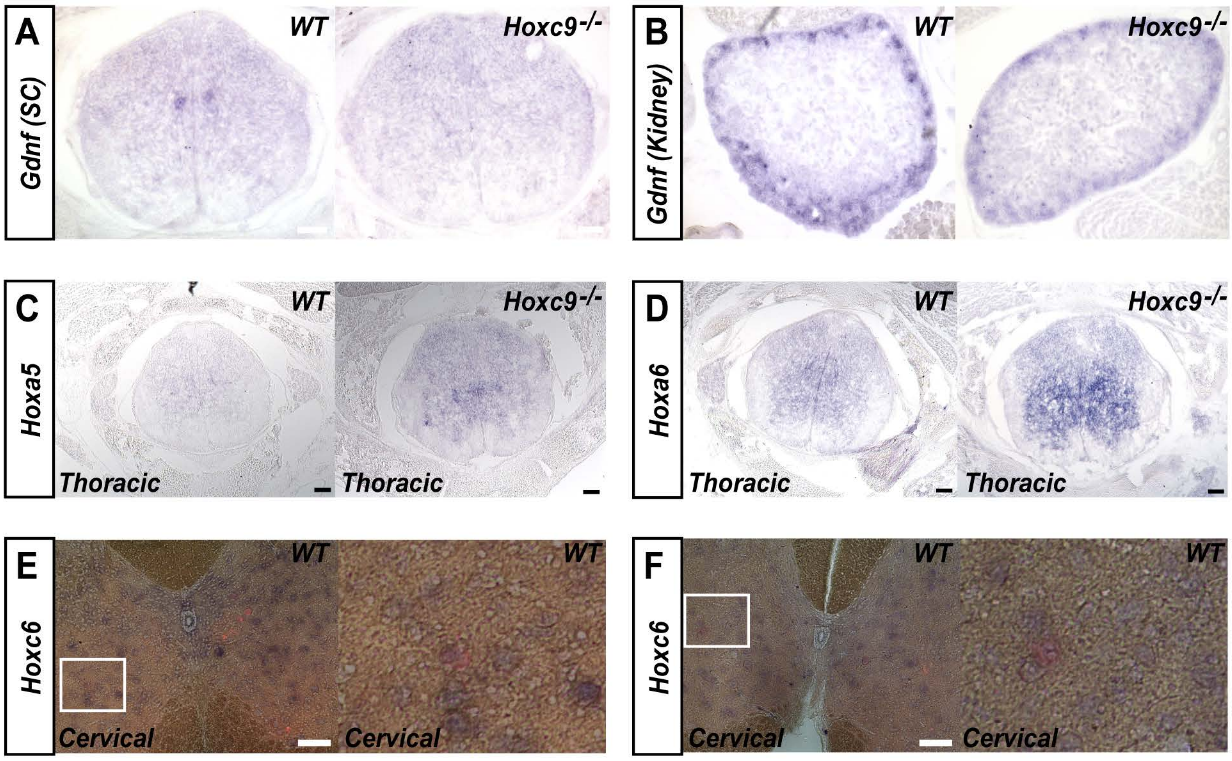

A
Gdnf (SC)
WT
Hoxc9-/-
B
Gdnf (Kidney)
WT
Hoxc9-/-
C
Hoxa5
WT
Hoxc9-/-
Thoracic
Thoracic
D
Hoxa6
WT
Hoxc9-/-
Thoracic
Thoracic
E
Hoxc6
WT
WT
Cervical
Cervical
F
Hoxc6
WT
WT
Cervical
Cervical
